# Antiviral efficacy of the SARS-CoV-2 XBB breakthrough infection sera against Omicron subvariants including EG.5

**DOI:** 10.1101/2023.08.08.552415

**Authors:** Yu Kaku, Yusuke Kosugi, Keiya Uriu, Jumpei Ito, Jin Kuramochi, Kenji Sadamasu, Kazuhisa Yoshimura, Hiroyuki Asakura, Mami Nagashima, The Genotype to Phenotype Japan (G2P-Japan) Consortium, Kei Sato

**Affiliations:** Division of Systems Virology, Department of Microbiology and Immunology, The Institute of Medical Science, The University of Tokyo, Tokyo, Japan; Graduate School of Medicine, The University of Tokyo, Tokyo, Japan; International Research Center for Infectious Diseases, The Institute of Medical Science, The University of Tokyo, Tokyo, Japan; Interpark Kuramochi Clinic, Utsunomiya, Japan; Department of Global Health Promotion, Tokyo Medical and Dental University, Tokyo, Japan; Tokyo Metropolitan Institute of Public Health, Tokyo, Japan; Graduate School of Frontier Sciences, The University of Tokyo, Kashiwa, Japan; International Vaccine Design Center, The Institute of Medical Science, The University of Tokyo, Tokyo, Japan; Collaboration Unit for Infection, Joint Research Center for Human Retrovirus infection, Kumamoto University, Kumamoto, Japan; CREST, Japan Science and Technology Agency, Kawaguchi, Japan

**Author notes:** Correspondence (Kei Sato). Contributed equally to this study (Yu Kaku, Yusuke Kosugi, Keiya Uriu, Jumpei Ito).

## Abstract

As of July 2023, EG.5.1 (a.k.a. XBB.1.9.2.5.1), a XBB subvariant bearing the S:Q52H and S:F456L substitutions, alongside the S:F486P substitution (Figure S1A), has rapidly spread in some countries. On July 19, 2023, the WHO classified EG.5 as a variant under monitoring. First, we showed that EG.5.1 exhibits a higher effective reproduction number compared with XBB.1.5, XBB.1.16, and its parental lineage (XBB.1.9.2), suggesting that EG.5.1 will spread globally and outcompete these XBB subvariants in the near future. We then addressed whether EG.5.1 evades from the antiviral effect of the humoral immunity induced by breakthrough infection (BTI) of XBB subvariants and performed a neutralization assay using XBB BTI sera. However, the 50% neutralization titer (NT50) of XBB BTI sera against EG.5.1 was comparable to those against XBB.1.5/1.9.2 and XBB.1.16. Moreover, the sensitivity of EG.5.1 to convalescent sera of XBB.1- and XBB.1.5-infected hamsters was similar to those of XBB.1.5/1.9 and XBB.1.16. These results suggest that the increased Re of EG.5.1 is attributed to neither increased infectivity nor immune evasion from XBB BTI, and the emergence and spread of EG.5 is driven by the other pressures. We previously demonstrated that Omicron BTI cannot efficiently induce antiviral humoral immunity against the variant infected. In fact, the NT50s of the BTI sera of Omicron BA.1, BA.2, and BA.5 against the variant infected were 3.0-, 2.2-, and 3.4-fold lower than that against the ancestral B.1.1 variant, respectively. However, strikingly, we found that the NT50 of the BTI sera of XBB1.5/1.9 and XBB.1.16 against the variant infected were 8.7- and 8.3-fold lower than that against the B.1.1 variant. These results suggest that XBB BTI cannot efficiently induce antiviral humoral immunity against XBB subvariants.

## Text

The SARS-CoV-2 XBB lineage is a recombinant Omicron lineage that emerged in the summer of 2022^1^. As of July 2023, the XBB sublineages, most notably those bearing the F486P substitution in the spike protein (S; S:F486P), such as XBB.1.5 and XBB.1.16, have rapidly spread and become predominant in the world (https://nextstrain.org/). As of July 2023, EG.5.1 (a.k.a. XBB.1.9.2.5.1), a XBB subvariant bearing the S:Q52H and S:F456L substitutions, alongside the S:F486P substitution (**Figure S1A**), has rapidly spread in some Asian and North American countries (**Figures S1B**). On July 19, 2023, the WHO classified EG.5 as a variant under monitoring.^2^ In fact, our analyses showed that EG.5.1 exhibits a higher effective reproduction number (Re) compared with XBB.1.5, XBB.1.16, and its parental lineage (XBB.1.9.2), suggesting that EG.5.1 will spread globally and outcompete these XBB subvariants in the near future (**Figure S1C**).

To assess the possibility that the enhanced infectivity of EG.5.1 contributes to its augmented Re, we prepared pseudoviruses with the S proteins of EG.5.1, XBB.1.5/1.9.2, and their derivatives. However, both S:Q52H and S:F456L did not increase pseudovirus infectivity, and the pseudovirus infectivity of EG.5.1 was significantly lower than that of its parental lineage (XBB.1.9.2) (**Figure S1D**). We then addressed whether EG.5.1 evades the antiviral effect of the humoral immunity induced by breakthrough infection (BTI) of XBB subvariants and performed a neutralization assay using XBB BTI sera. However, the 50% neutralization titer (NT50) of XBB BTI sera against EG.5.1 was comparable to those against XBB.1.5/1.9.2 and XBB.1.16 (**Figure S1E**). Moreover, the sensitivity of EG.5.1 to convalescent sera of XBB.1- and XBB.1.5-infected hamsters was similar to those of XBB.1.5/1.9 and XBB.1.16 (**Figure S1F**). These results suggest that the increased Re of EG.5.1 is attributed to neither increased infectivity nor immune evasion from XBB BTI, and the emergence and spread of EG.5 is driven by the other pressures.

We previously demonstrated that Omicron BTI cannot efficiently induce antiviral humoral immunity against the variant infected.^1, 3–6^ In fact, the NT50s of the BTI sera of Omicron BA.1, BA.2, and BA.5 against the variant infected were 3.0-, 2.2-, and 3.4-fold lower than that against the ancestral B.1.1 variant, respectively (**Figure S1G**). Strikingly, however, we found that the NT50 of the BTI sera of XBB1.5/1.9.2 and XBB.1.16 against the variant infected were 8.7- and 8.3-fold lower than that against the B.1.1 variant (**Figure S1E**). These results suggest that XBB BTI cannot efficiently induce antiviral humoral immunity against XBB subvariants.

## Grants

Supported in part by AMED SCARDA Japan Initiative for World-leading Vaccine Research and Development Centers “UTOPIA” (JP223fa627001, to Kei Sato), AMED SCARDA Program on R&D of new generation vaccine including new modality application (JP223fa727002, to Kei Sato); AMED Research Program on Emerging and Re-emerging Infectious Diseases (JP22fk0108146, to Kei Sato; JP21fk0108494 to G2P-Japan Consortium and Kei Sato; JP21fk0108425, to Kei Sato; JP21fk0108432, to Kei Sato; JP22fk0108511, to G2P-Japan Consortium and Kei Sato; JP22fk0108516, to Kei Sato; JP22fk0108506, to Kei Sato); AMED Research Program on HIV/AIDS (JP22fk0410039, to Kei Sato); JST PRESTO (JPMJPR22R1, to Jumpei Ito); JST CREST (JPMJCR20H4, to Kei Sato); JSPS KAKENHI Grant-in-Aid for Early-Career Scientists (23K14526, to Jumpei Ito); JSPS Core-to-Core Program (A. Advanced Research Networks) (JPJSCCA20190008, Kei Sato); JSPS Research Fellow DC2 (22J11578, to Keiya Uriu); JSPS Research Fellow DC1 (23KJ0710, to Yusuke Kosugi); The Tokyo Biochemical Research Foundation (to Kei Sato); The Mitsubishi Foundation (to Kei Sato).

## Declaration of interest

J.I. has consulting fees and honoraria for lectures from Takeda Pharmaceutical Co. Ltd. K.S. has consulting fees from Moderna Japan Co., Ltd. and Takeda Pharmaceutical Co. Ltd. and honoraria for lectures from Gilead Sciences, Inc., Moderna Japan Co., Ltd., and Shionogi & Co., Ltd. The other authors declare no competing interests. All authors have submitted the ICMJE Form for Disclosure of Potential Conflicts of Interest. Conflicts that the editors consider relevant to the content of the manuscript have been disclosed.

Conflict of interest: The authors declare that no competing interests exist.

## Supplementary Appendix

### Supplementary Discussion

On June 15, 2023, the FDA’s Vaccines and Related Biological Products Advisory Committee issued a recommendation for the use of the monovalent XBB.1.5 vaccine for the purpose of booster vaccination in the autumn of 2023.^7^ Because booster vaccination with XBB.1.5-containing vaccines induces >10-fold anti-XBB neutralizing antibodies,^8^ the monovalent XBB.1.5 vaccine may potentially induce anti-XBB humoral immunity. However, here we showed that the immunogenicity of XBB subvariants is lower than that of previous Omicron subvariants (**Figure S1E and S1G**). Therefore, the immune activation induced by XBB BTI and further XBB vaccination may be less effective than expected. It should be noted that our results presented here do not imply ineffectiveness of XBB-containing vaccines; however, our data suggest that XBB BTI does not sufficiently induce anti-XBB humoral immunity.

## Materials and Methods

### Ethics statement

All protocols involving specimens from human subjects recruited at Interpark Kuramochi Clinic was reviewed and approved by the Institutional Review Board of Interpark Kuramochi Clinic (approval ID: G2021-004). All human subjects provided written informed consent. All protocols for the use of human specimens were reviewed and approved by the Institutional Review Boards of The Institute of Medical Science, The University of Tokyo (approval IDs: 2021-1-0416 and 2021-18-0617).

### Human serum collection

Convalescent sera were collected from fully vaccinated individuals who had been infected with BA.1 (thirteen 2-dose vaccinated; time interval between the last vaccination and infection, 71– 251 days; 7–27 days after testing. n=13; average age: 43 years, range: 20–65 years, 46.2% male) (**Figure S1G**), BA.2 (nine 2-dose vaccinated and four 3-dose vaccinated; time interval between the last vaccination and infection, 4–299 days; 11–61 days after testing. n=13 in total; average age: 45 years, range: 24–82 years, 62% male) (**Figure S1G**), BA.5 (one 2-dose vaccinated, thirteen 3-dose vaccinated and one 4-dose vaccinated; time interval between the last vaccination and infection, 66–310 days; 10–23 days after testing. n=15 in total; average age: 55 years, range: 25–73 years, 47% male) (**Figure S1G**), and XBB sublineages [XBB.1.5 (one 3-dose vaccinated; time interval between the last vaccination and infection, 413 days; 38 days after testing. n=1 in total; average age: 55 years, range: 55 years, 100% male), XBB.1.9 (one 3-dose vaccinated, one 4-dose vaccinated and one 5-dose vaccinated; time interval between the last vaccination and infection, 176–246 days; 6–18 days after testing. n=3 in total; average age: 53.7 years, range: 32–88 years, 33.3% male), and XBB.1.16 (one 2-dose vaccinated, two 3-dose vaccinated and one 4-dose vaccinated; time interval between the last vaccination and infection, 223–569 days; 7–15 days after testing. n=4 in total; average age: 38.8 years, range: 18–65 years, 100% male)] (**Figure S1E**). The SARS-CoV-2 variants were identified as previously described.^1–3^ Sera were inactivated at 56°C for 30 minutes and stored at –80°C until use. The details of the convalescent sera are summarized in **Table S1**.

### Hamster serum collection

Animal experiments were performed as previously described.^1–14^ Briefly, 4-week male Syrian hamsters purchased from Japan SLC Inc. (Shizuoka, Japan) were inoculated with 10,000 50% tissue culture infectious dose (TCID50) of XBB.1 or XBB.1.5 via the intranasal route under anesthesia. The anesthetics used for viral infection were injected into muscles as a mixture of 0.15 mg/kg medetomidine hydrochloride (Domitor®, Nippon Zenyaku Kogyo), 2.0 mg/kg midazolam (Dormicum®, Fujifilm Wako, Cat# 135-13791) and 2.5 mg/kg butorphanol (Vetorphale®, Meiji Seika Pharma) or 0.15 mg/kg medetomidine hydrochloride, 4.0 mg/kg alphaxaone (Alfaxan®, Jurox) and 2.5 mg/kg butorphanol at 16 days postinfection.

### Epidemic dynamics analysis

In the present study, we analyzed the viral genomic surveillance data deposited in the GISAID database (https://www.gisaid.org/; downloaded on July 13, 2023). We used the data from April 1, 2023 in this analysis. We excluded the sequence records with the following features: i) a lack of collection date information; ii) sampling in animals other than humans; iii) sampling by quarantine; iv) without the PANGO lineage information; and v) having >2% undetermined (N) nucleotide sequences. In the downstream analysis, we only used sequences for PANGO lineages with >50 sequences in the dataset. We modeled the epidemic dynamics of viral lineages in six countries (China, South Korea, USA, Japan, Singapore, and Canada), where >50 sequences of XBB.1.5, XBB.1.16, and EG.5.1 were detected. We counted the daily frequency of each viral lineage. Subsequently, epidemic dynamics and relative Re value for each viral lineage were estimated according to the Bayesian multinomial logistic model, described in our previous study.^2^ Briefly, we estimated the logistic slope parameter β_*l*_ for each viral lineage using the model and then calculated relative Re for each lineage 𝑟_*l*_ as 𝑟_*l*_ = 𝑒𝑥𝑝(γβ_*l*_) where γ is the average viral generation time (2.1 days) (http://sonorouschocolate.com/covid19/index.php?title=Estimating_Generation_Time_Of_Omicron). For parameter estimation, the intercept and slope parameters of XBB.1.5 were fixed at 0. Consequently, the relative Re of XBB.1.5 was fixed at 1, and those of the other lineages were estimated relative to that of XBB.1. Parameter estimation was performed via the MCMC approach implemented in CmdStan v2.31.0 (https://mc-stan.org) with CmdStanr v0.5.3 (https://mc-stan.org/cmdstanr/). Four independent MCMC chains were run with 1,000 and 4,000 steps in the warmup and sampling iterations, respectively. We confirmed that all estimated parameters showed <1.05 R-hat convergence diagnostic values and >200 effective sampling size values, indicating that the MCMC runs were successfully convergent. Information on the estimated parameters is summarized in **Table S2**. In **Figures S1B and S1C**, results for XBB.1.5, XBB.1.9.2, XBB.1.16, and EG.5.1 are shown.

### Plasmid construction

Plasmids expressing the SARS-CoV-2 spike proteins of the ancestral B.1.1 (D614G-bearing virus), Omicron BA.1, BA.2, BA.5, BQ.1.1, XBB.1, XBB.1.5/1.9.2 and XBB.1.16 were prepared in our previous studies.^1, 2, 4,15–19^ Note that the S proteins of XBB.1.5 and XBB.1.9.2 are identical (**Figure S1A**). Plasmids expressing the spike protein of EG.5.1 and its derivative were generated by site-directed overlap extension PCR using pC-SARS2-S XBB.1.5^18^ as the template and the primers listed in **Table S3**. The resulting PCR fragment was subcloned into the KpnI-NotI site of the pCAGGS vector^20^ using In-Fusion HD Cloning Kit (Takara, Cat# Z9650N). Nucleotide sequences were determined by DNA sequencing services (Eurofins), and the sequence data were analyzed by SnapGene software v6.1.1 (www.snapgene.com).

### Cell culture

The Lenti-X 293T cell line (Takara, Cat# 632180) and HOS-ACE2/TMPRSS2 cells (kindly provided by Dr. Kenzo Tokunaga), a derivative of HOS cells (a human osteosarcoma cell line; ATCC CRL-1543) stably expressing human ACE2 and TMPRSS2,^7, 21^ were maintained in Dulbecco’s modified Eagle’s medium (DMEM) (high glucose) (Wako, Cat# 044-29765) containing 10% fetal bovine serum (Sigma-Aldrich Cat# 172012-500ML), 100 units penicillin and 100 ug/ml streptomycin (Sigma-Aldrich, Cat# P4333-100ML).

### Pseudovirus preparation

Pseudoviruses were prepared as previously described.^1–5, 7–11, 13, 15–19, 22^ Briefly, lentivirus (HIV-1)-based, luciferase-expressing reporter viruses were pseudotyped with the SARS-CoV-2 spikes. HEK293T cells (2 × 10^6^ cells) were cotransfected with 1 μg psPAX2-IN/HiBiT,^23^ 1 μg pWPI-Luc2,^23^ and 500 ng plasmids expressing parental S or its derivatives using TransIT-293 transfection reagent (Mirus, Cat# MIR2704) according to the manufacturer’s protocol. Two days post transfection, the culture supernatants were harvested and centrifuged. The amount of pseudoviruses prepared was quantified by the HiBiT assay using Nano Glo HiBiT lytic detection system (Promega,Cat# N3040) as previously described.^1–5, 7–11, 13, 15–19, 22^ To measure viral infectivity, the same amount of pseudoviruses (normalized to the HiBiT value, which indicates the amount of p24 HIV-1 antigen) was inoculated into HOS-ACE2/TMPRSS2 cells. At two days postinfection, the infected cells were lysed with a Bright-Glo luciferase assay system (Promega, Cat# E2620), and the luminescent signal was measured using a GloMax explorer multimode microplate reader 3500 (Promega). The pseudoviruses were stored at –80°C until use.

### Neutralization assay

Neutralization assays were performed as previously described.^1–5, 7, 8, 10, 11, 13, 16–19^ The SARS-CoV-2 spike pseudoviruses (counting ∼100,000 relative light units) were incubated with serially diluted (120-fold to 87,480-fold dilution at the final concentration) heat-inactivated sera at 37°C for 1 hour. Pseudoviruses without sera were included as controls. Then, 20 μl mixture of pseudovirus and serum was added to HOS-ACE2/TMPRSS2 cells (10,000 cells/100 μl) in a 96-well white plate. Two days post infection, the infected cells were lysed with a Bright-Glo luciferase assay system (Promega, Cat# E2620), and the luminescent signal was measured using a GloMax explorer multimode microplate reader 3500 (Promega). The assay of each serum sample was performed in triplicate, and the 50% neutralization titer (NT50) was calculated using Prism 9 (GraphPad Software).

### Data availability

Dataset used in the epidemic dynamics analysis in this study is available from the GISAID database (https://www.gisaid.org; EPI_SET_230725pv). The GISAID supplemental tables for EPI_SET_230725pv is available in the GitHub repository (https://github.com/TheSatoLab/EG.5.1_short).

## Supplementary figure

**Table S1.** Human sera used in this study

| SARS-CoV-2 infected | Donor ID | Sex | Age | vaccine<br>(1st vaccination) | Date of 1st vaccination<br>(YYYY-MM-DD) | vaccine<br>(2nd vaccination) | Date of 2nd vaccination<br>(YYYY-MM-DD) | vaccine<br>(3rd vaccination) | Date of 3rd vaccination<br>(YYYY-MM-DD) | vaccine<br>(4th vaccination) | Date of 4th vaccination<br>(YYYY-MM-DD) | vaccine<br>(5th vaccination) | Date of 5th vaccination<br>(YYYY-MM-DD) | Date of test<br>(YYYY-MM-DD) | Date of sampling<br>(YYYY-MM-DD) | Prior<br>infection? |
| --- | --- | --- | --- | --- | --- | --- | --- | --- | --- | --- | --- | --- | --- | --- | --- | --- |
| BA.1 | P264 | Male | 60 | BNT162b2 | 2021-08-27 | BNT162b2 | 2021-09-18 |  |  |  |  |  |  | 2022-01-03 | 2022-01-29 | No |
| BA.1 | P265 | Female | 57 | BNT162b2 | 2021-10-04 | BNT162b2 | 2021-10-25 |  |  |  |  |  |  | 2022-01-04 | 2022-01-28 | No |
| BA.1 | P274 | Female | 20 | mRNA-1273 | 2021-09-11 | mRNA-1273 | 2021-10-09 |  |  |  |  |  |  | 2022-01-15 | 2022-01-29 | No |
| BA.1 | P276 | Male | 44 | BNT162b2 | 2021-08-06 | BNT162b2 | 2021-08-22 |  |  |  |  |  |  | 2022-01-20 | 2022-01-30 | No |
| BA.1 | P279 | Male | 61 | BNT162b2 | 2021-07-03 | BNT162b2 | 2021-07-24 |  |  |  |  |  |  | 2022-01-18 | 2022-02-05 | No |
| BA.1 | P282 | Male | 56 | BNT162b2 | 2021-09-18 | BNT162b2 | 2021-10-12 |  |  |  |  |  |  | 2022-01-22 | 2022-02-06 | No |
| BA.1 | P285 | Female | 65 | BNT162b2 | 2022-01-07 | BNT162b2 | 2022-01-28 |  |  |  |  |  |  | 2022-01-23 | 2022-02-06 | No |
| BA.1 | P295 | Female | 41 | BNT162b2 | 2021-08-07 | BNT162b2 | 2021-08-28 |  |  |  |  |  |  | 2022-01-25 | 2022-02-06 | No |
| BA.1 | 2112 | Male | 44 | BNT162b2 | 2021-04-21 | BNT162b2 | 2021-05-12 |  |  |  |  |  |  | 2022-01-18 | 2022-01-25 | No |
| BA.1 | 1880 | Female | 25 | BNT162b2 | 2021-09-09 | BNT162b2 | 2021-09-30 |  |  |  |  |  |  | 2022-01-07 | 2022-02-03 | No |
| BA.1 | 2187 | Male | 32 | mRNA-1273 | 2021-07-09 | mRNA-1273 | 2021-08-06 |  |  |  |  |  |  | 2022-01-20 | 2022-02-05 | No |
| BA.1 | 2137 | Female | 32 | mRNA-1273 | 2021-07-09 | mRNA-1273 | 2021-08-06 |  |  |  |  |  |  | 2022-01-19 | 2022-02-05 | No |
| BA.1 | 2550 | Female | 24 | BNT162b2 | 2021-10-12 | BNT162b2 | 2021-11-02 |  |  |  |  |  |  | 2022-01-25 | 2022-02-05 | No |
| BA.2 | P378 | Male | 43 | BNT162b2 | 2021-10-10 | BNT162b2 | 2021-10-31 | BNT162b2 |  |  |  |  |  | 2022-03-28 | 2022-04-10 | No |
| BA.2 | P398 | Male | 48 | BNT162b2 | 2021-09-18 | BNT162b2 | 2021-10-09 | BNT162b2 | 2022-04-09 |  |  |  |  | 2022-04-13 | 2022-04-30 | No |
| BA.2 | P407 | Male | 29 | mRNA-1273 | 2021-09-13 | mRNA-1273 | 2021/10/11 | mRNA-1273 |  |  |  |  |  | 2022-05-01 | 2022-05-12 | No |
| BA.2 | P401 | Male | 35 | BNT162b2 | 2021-09-09 | BNT162b2 | 2021-09-30 | BNT162b2 |  |  |  |  |  | 2022-04-22 | 2022-05-05 | No |
| BA.2 | P412 | Female | 82 | BNT162b2 | 2021-06-11 | BNT162b2 | 2021-07-09 | BNT162b2 |  |  |  |  |  | 2022-05-04 | 2022-05-26 | No |
| BA.2 | 6449 | Male | 43 | BNT162b2 | 2021-08-13 | BNT162b2 | 2021-09-11 | BNT162b2 |  |  |  |  |  | 2022-04-03 | 2022-04-23 | No |
| BA.2 | 6355 | Male | 50 | NA | 2021-04-28 | NA | 2021-05-19 | NA | 2022-01-19 |  |  |  |  | 2022-04-02 | 2022-04-20 | No |
| BA.2 | 6547 | Male | 54 | BNT162b2 | 2021-08-25 | BNT162b2 | 2021-09-15 | BNT162b2 |  |  |  |  |  | 2022-04-06 | 2022-04-22 | No |
| BA.2 | 7951 | Female | 71 | BNT162b2 | 2021-06-20 | BNT162b2 | 2021-07-16 | mRNA-1273 | 2022-02-16 |  |  |  |  | 2022-04-25 | 2022-05-12 | No |
| BA.2 | 8645 | Female | 41 | BNT162b2 | 2021-05-23 | BNT162b2 | 2021-06-13 | BNT162b2 | 2022-01-20 |  |  |  |  | 2022-05-07 | 2022-05-20 | No |
| BA.2 | 8682 | Female | 25 | BNT162b2 | 2021-09-03 | BNT162b2 | 2021-09-27 | BNT162b2 |  |  |  |  |  | 2022-05-08 | 2022-05-24 | No |
| BA.2 | 5949 | Male | 24 | mRNA-1273 | 2021-08-05 | mRNA-1273 | 2021-09-02 | mRNA-1273 |  |  |  |  |  | 2022-03-22 | 2022-05-22 | No |
| BA.2 | 8796 | Female | 34 | BNT162b2 | 2021-09-26 | BNT162b2 | 2021-10-17 | BNT162b2 |  |  |  |  |  | 2022-05-10 | 2022-06-05 | No |
| BA.5 | P427 | Female | 49 | NA | 2021-07-30 | NA | 2021-08-25 | NA | 2022-03-18 |  |  |  |  | 2022-07-06 | 2022-07-25 | No |
| BA.5 | P439 | Female | 73 | BNT162b2 | 2021-06-19 | BNT162b2 | 2021-07-20 | mRNA-1273 | 2022-02-04 |  |  |  |  | 2022-07-23 | 2022-08-08 | No |
| BA.5 | P451 | Female | 55 | BNT162b2 | 2021-04-26 | BNT162b2 | 2021-05-20 | BNT162b2 | 2022-01-18 |  |  |  |  | 2022-07-29 | 2022-08-12 | No |
| BA.5 | P456 | Male | 44 | BNT162b2 | 2021-08-11 | BNT162b2 | 2021-09-01 | mRNA-1273 | 2022-03-13 |  |  |  |  | 2022-08-04 | 2022-08-14 | No |
| BA.5 | P464 | Male | 63 | BNT162b2 | 2021-08-08 | BNT162b2 | 2021-08-29 | mRNA-1273 | 2022-04-07 |  |  |  |  | 2022-08-08 | 2022-08-19 | No |
| BA.5 | 9341 | Male | 56 | BNT162b2 | 2021-08-10 | BNT162b2 | 2021-08-31 | BNT162b2 | 2022-03-18 |  |  |  |  | 2022-06-12 | 2022-06-30 | No |
| BA.5 | 9584 | Male | 55 | BNT162b2 | 2021-07-14 | BNT162b2 | 2021-08-05 | BNT162b2 | 2022-03-22 |  |  |  |  | 2022-07-08 | 2022-07-25 | No |
| BA.5 | 11318 | Female | 51 | BNT162b2 | 2021-09-01 | BNT162b2 | 2021-09-22 | mRNA-1273 | 2022-05-19 |  |  |  |  | 2022-07-24 | 2022-08-05 | No |
| BA.5 | 23S-08 | Male | 25 | BNT162b2 | 2021-04-27 | BNT162b2 | 2021-05-18 | BNT162b2 | 2022-01-11 |  |  |  |  | 2022-07-23 | 2022-08-08 | No |
| BA.5 | 10826 | Male | 63 | BNT162b2 | 2021-07-27 | BNT162b2 | 2021-08-17 | BNT162b2 | 2022-03-04 | mRNA-1273 | 2022-08-09 |  |  | 2022-07-21 | 2022-08-11 | No |
| BA.5 | 11079 | Female | 65 | mRNA-1273 | 2021-07-08 | mRNA-1273 | 2021-08-05 | mRNA-1273 | 2022-03-17 |  |  |  |  | 2022-07-23 | 2022-08-11 | No |
| BA.5 | 14847 | Female | 70 | BNT162b2 | 2021-07-13 | BNT162b2 | 2021-08-20 | mRNA-1273 | 2022-03-08 |  |  |  |  | 2022-08-13 | 2022-08-25 | No |
| BA.5 | 13180 | Female | 63 | BNT162b2 | 2021-07-16 | BNT162b2 | 2021-08-06 | mRNA-1273 | 2022-03-08 |  |  |  |  | 2022-08-04 | 2022-08-25 | No |
| BA.5 | 12912 | Male | 64 | mRNA-1273 | 2021-09-02 | mRNA-1273 | 2021-09-30 | mRNA-1273 | 2021-04-01 |  |  |  |  | 2022-08-02 | 2022-08-25 | No |
| BA.5 | 14956 | Female | 33 | BNT162b2 | 2021-09-06 | BNT162b2 | 2021-10-07 |  |  |  |  |  |  | 2022-08-13 | 2022-08-28 | No |
| XBB.1.5 | 36708 | Male | 55 | BNT162b2 | 2021-08-07 | BNT162b2 | 2021-08-27 | BNT162b2 | 2022-04-14 |  |  |  |  | 2023-06-01 | 2023-07-09 | No |
| XBB.1.9 | P592 | Female | 32 | NA | NA | NA | NA | BNT162b2 | 2022-10-26 |  |  |  |  | 2023-05-06 | 2023-05-16 | No |
| XBB.1.9 | P595 | Female | 88 | BNT162b2 | 2021-05-18 | BNT162b2 | 2021-06-08 | mRNA-1273 | 2022-02-04 | mRNA-1273 | 2022-07-19 | bivalent, Pfizer-BioNTech | 2022-11-18 | 2023-05-13 | 2023-05-19 | No |
| XBB.1.9 | KS-230718 | Male | 41 | BNT162b2 | 2021-06-17 | BNT162b2 | 2021-07-07 | mRNA-1273 | 2022-03-28 | mRNA-1273.222 | 2022-10-27 |  |  | 2023-06-30 | 2023-07-18 | No |
| XBB.1.16 | P601 | Male | 38 | BNT162b2 | 2021-11-05 | BNT162b2 | 2021-11-26 | BNT162b2 | 2022-05-27 |  |  |  |  | 2023-07-06 | 2023-07-14 | No |
| XBB.1.16 | P593 | Male | 65 | BNT162b2 | 2021-08-02 | BNT162b2 | 2021-08-30 | BNT162b2 | 2022-04-03 |  |  |  |  | 2023-05-09 | 2023-05-16 | No |
| XBB.1.16 | P594 | Male | 18 | BNT162b2 | 2021-09-06 | BNT162b2 | 2021-10-15 |  |  |  |  |  |  | 2023-05-07 | 2023-05-19 | No |
| XBB.1.16 | JI-230718 | Male | 34 | mRNA-1273 | 2021-07-19 | mRNA-1273 | 2021-08-20 | mRNA-1273 | 2022-05-13 | mRNA-1273.222 | 2022-11-22 |  |  | 2023-07-03 | 2023-07-18 | Yes |

NA, not applicable.

**Table S2.** Estimated relative Re and epidemic dynamics modeling parameters of the representative SARS-CoV-2 Omicron sublineages spreading.

| Country | PANGO lineage | Posterior mean | Posterior 2.5 percentile | Posterior 97.5 percentile | R-hat value | Effective sampling size (ESS_bulk) | Effective sampling size (ess_tail) |
| --- | --- | --- | --- | --- | --- | --- | --- |
| Canada | EG.5.1 | 1.194 | 1.156 | 1.235 | 1.000 | 28020.167 | 11760.151 |
| Canada | XBB.1.5.10 | 1.130 | 1.098 | 1.163 | 1.000 | 28852.170 | 11060.454 |
| Canada | XBB.1.5.77 | 1.129 | 1.099 | 1.160 | 1.000 | 26316.643 | 11847.772 |
| Canada | FL.2 | 1.117 | 1.087 | 1.148 | 1.000 | 23850.488 | 12222.184 |
| Canada | FL.4 | 1.106 | 1.093 | 1.118 | 1.000 | 21557.407 | 11294.275 |
| Canada | EG.5.2 | 1.098 | 1.070 | 1.127 | 1.001 | 22065.019 | 11225.833 |
| Canada | XBB.1.9.2 | 1.093 | 1.077 | 1.110 | 1.000 | 24401.962 | 11272.720 |
| Canada | XBB.1.16 | 1.092 | 1.085 | 1.100 | 1.000 | 16188.077 | 12184.423 |
| Canada | XBB.1.16.3 | 1.091 | 1.065 | 1.117 | 1.000 | 21908.154 | 12291.440 |
| Canada | XBB.1.16.1 | 1.087 | 1.073 | 1.101 | 1.000 | 22902.095 | 11224.878 |
| Canada | FL.5 | 1.077 | 1.061 | 1.095 | 1.000 | 24573.253 | 11876.335 |
| Canada | EG.1 | 1.076 | 1.059 | 1.093 | 1.000 | 25130.931 | 11995.833 |
| Canada | FE.1.1.1 | 1.073 | 1.053 | 1.094 | 1.000 | 24534.713 | 12577.649 |
| Canada | FD.1.1 | 1.071 | 1.063 | 1.078 | 1.000 | 19787.553 | 12077.306 |
| Canada | XBB.1.16.2 | 1.070 | 1.052 | 1.088 | 1.001 | 23073.775 | 12690.091 |
| Canada | XBB.1.5.37 | 1.069 | 1.047 | 1.092 | 1.000 | 27311.861 | 11906.712 |
| Canada | XBB.1.9.1 | 1.067 | 1.058 | 1.076 | 1.000 | 19615.125 | 12280.830 |
| Canada | XBB.2.3.2 | 1.066 | 1.044 | 1.087 | 1.000 | 25803.316 | 12386.694 |
| Canada | FL.13 | 1.064 | 1.043 | 1.087 | 1.000 | 26894.810 | 11857.820 |
| Canada | XBB.2.3.3 | 1.059 | 1.034 | 1.083 | 1.000 | 22087.655 | 11522.770 |
| Canada | XBB.2.3 | 1.046 | 1.029 | 1.063 | 1.001 | 25501.988 | 12326.880 |
| Canada | XBB.1.5.62 | 1.039 | 1.010 | 1.068 | 1.001 | 28943.932 | 11380.550 |
| Canada | XBB.1.5.15 | 1.034 | 1.013 | 1.054 | 1.000 | 25653.395 | 11745.767 |
| Canada | XBB.1.5.4 | 1.026 | 1.002 | 1.049 | 1.000 | 25794.898 | 11808.858 |
| Canada | XBB.1.5.13 | 1.023 | 1.008 | 1.038 | 1.001 | 26031.661 | 12118.199 |
| Canada | XBB.1.5.49 | 1.014 | 1.000 | 1.027 | 1.000 | 23714.177 | 12657.722 |
| Canada | XBB.1.5.17 | 0.999 | 0.979 | 1.019 | 1.000 | 28516.422 | 11615.119 |
| Canada | CH.1.1 | 0.998 | 0.972 | 1.025 | 1.000 | 30231.545 | 12166.557 |
| Canada | DV.1.1 | 0.995 | 0.979 | 1.010 | 1.000 | 25337.707 | 12063.520 |
| Canada | XBB.1.5.1 | 0.993 | 0.977 | 1.009 | 1.000 | 23550.112 | 12375.932 |
| Canada | XBB.1.5.50 | 0.993 | 0.975 | 1.010 | 1.000 | 29322.905 | 12455.737 |
| Canada | XBB.1.5.19 | 0.992 | 0.962 | 1.022 | 1.000 | 27375.832 | 12903.215 |
| Canada | XBB.1.5.12 | 0.990 | 0.959 | 1.020 | 1.000 | 25564.766 | 12585.821 |
| Canada | XBB.1.5.2 | 0.983 | 0.963 | 1.001 | 1.001 | 24836.675 | 11648.676 |
| Canada | XBB.1.5.31 | 0.982 | 0.957 | 1.006 | 1.000 | 25229.769 | 12823.027 |
| Canada | XBB.1.5.47 | 0.979 | 0.955 | 1.002 | 1.000 | 25909.971 | 11770.983 |
| Canada | XBB.1.5.32 | 0.977 | 0.965 | 0.988 | 1.000 | 26107.411 | 12173.604 |
| Canada | XBB.1.5.7 | 0.976 | 0.956 | 0.996 | 1.000 | 25225.149 | 12498.934 |
| Canada | XBB.1.5.20 | 0.970 | 0.953 | 0.988 | 1.000 | 27428.240 | 12096.116 |
| Canada | FT.1 | 0.945 | 0.925 | 0.965 | 1.000 | 26792.440 | 11491.241 |
| Canada | XBB.1.5.69 | 0.934 | 0.904 | 0.963 | 1.000 | 25785.623 | 12281.238 |
| Canada | BQ.1.1 | 0.857 | 0.822 | 0.889 | 1.000 | 28919.825 | 12538.675 |
| China | EG.5.1 | 1.198 | 1.182 | 1.214 | 1.001 | 3394.461 | 6749.947 |
| China | FU.1 | 1.112 | 1.099 | 1.124 | 1.001 | 2866.909 | 5047.291 |
| China | FY.3 | 1.107 | 1.092 | 1.123 | 1.001 | 3766.592 | 7191.155 |
| China | XBB.1.16.2 | 1.106 | 1.085 | 1.127 | 1.001 | 5505.439 | 9232.494 |
| China | FR.1.1 | 1.105 | 1.087 | 1.122 | 1.001 | 4762.218 | 8375.087 |
| China | FL.16 | 1.097 | 1.077 | 1.116 | 1.001 | 5251.397 | 8838.607 |
| China | XBB.1.42 | 1.096 | 1.073 | 1.118 | 1.001 | 6113.867 | 10367.651 |
| China | FY.3.1 | 1.096 | 1.080 | 1.112 | 1.001 | 4022.764 | 8396.949 |
| China | XBB.2.3.2 | 1.089 | 1.056 | 1.123 | 1.000 | 9933.624 | 11902.980 |
| China | FE.1.1 | 1.087 | 1.067 | 1.108 | 1.001 | 6257.855 | 8976.966 |
| China | EG.2 | 1.085 | 1.059 | 1.111 | 1.000 | 7397.494 | 10101.266 |
| China | FR.1.3 | 1.081 | 1.052 | 1.110 | 1.000 | 9120.002 | 10638.645 |
| China | GF.1 | 1.074 | 1.058 | 1.090 | 1.001 | 4047.622 | 7973.134 |
| China | FL.4 | 1.071 | 1.058 | 1.084 | 1.001 | 3208.886 | 5673.191 |
| China | XBB.1.16 | 1.064 | 1.052 | 1.077 | 1.001 | 3016.906 | 5799.148 |
| China | XBB.1.16.1 | 1.063 | 1.043 | 1.083 | 1.000 | 5912.777 | 9298.074 |
| China | FL.2.3 | 1.060 | 1.046 | 1.074 | 1.001 | 3452.619 | 6923.746 |
| China | XBB.1.9.1 | 1.058 | 1.044 | 1.072 | 1.001 | 3769.872 | 7041.165 |
| China | XBB.1.5.15 | 1.051 | 1.025 | 1.076 | 1.001 | 7875.051 | 10508.907 |
| China | FL.2 | 1.048 | 1.034 | 1.062 | 1.001 | 3892.654 | 7507.503 |
| China | FL.13 | 1.046 | 1.012 | 1.081 | 1.000 | 10088.378 | 10666.178 |
| China | FL.5 | 1.037 | 1.013 | 1.061 | 1.000 | 7368.311 | 10809.312 |
| China | GR.1 | 1.036 | 1.000 | 1.072 | 1.000 | 10610.031 | 11941.475 |
| China | FL.13.1 | 1.030 | 1.008 | 1.051 | 1.001 | 6230.363 | 9594.446 |
| China | XBB.1.9.2 | 1.026 | 1.003 | 1.051 | 1.001 | 7180.414 | 9327.159 |
| China | XBB.1.5.24 | 1.023 | 0.991 | 1.056 | 1.000 | 9366.787 | 11048.381 |
| China | XBB.1.44.1 | 1.013 | 0.982 | 1.043 | 1.000 | 10452.977 | 11376.953 |
| China | EG.4 | 1.001 | 0.970 | 1.032 | 1.000 | 10817.353 | 12100.357 |
| China | FR.1 | 0.986 | 0.954 | 1.018 | 1.000 | 9206.885 | 11015.016 |
| China | BN.1.2 | 0.985 | 0.951 | 1.019 | 1.001 | 11669.478 | 11050.626 |
| China | FL.4.2 | 0.982 | 0.951 | 1.012 | 1.000 | 10452.629 | 12414.800 |
| China | XBB.1.19.1 | 0.928 | 0.905 | 0.952 | 1.000 | 7664.880 | 10294.506 |
| China | BF.7.14.5 | 0.824 | 0.785 | 0.862 | 1.000 | 12552.157 | 12083.006 |
| China | DY.1 | 0.819 | 0.791 | 0.846 | 1.000 | 11315.141 | 11851.783 |
| China | BF.7.14.1 | 0.810 | 0.774 | 0.847 | 1.000 | 13021.484 | 12119.375 |
| China | DY.2 | 0.799 | 0.782 | 0.816 | 1.001 | 6947.947 | 10625.961 |
| China | DY.4 | 0.795 | 0.771 | 0.820 | 1.000 | 11117.544 | 12424.806 |
| China | BF.7.14 | 0.792 | 0.777 | 0.808 | 1.001 | 6442.040 | 9737.975 |
| China | BA.5.2.48 | 0.785 | 0.759 | 0.810 | 1.000 | 10194.251 | 11262.329 |
| Japan | XBB.1.5.5 | 1.185 | 1.149 | 1.224 | 1.000 | 20979.776 | 11252.480 |
| Japan | EG.5.1 | 1.154 | 1.127 | 1.183 | 1.000 | 16970.482 | 12000.980 |
| Japan | FK.1.3 | 1.092 | 1.063 | 1.121 | 1.000 | 19113.664 | 12266.279 |
| Japan | XBB.1.16.2 | 1.074 | 1.054 | 1.094 | 1.000 | 14570.489 | 11481.217 |
| Japan | XBB.2.3.2 | 1.060 | 1.042 | 1.078 | 1.000 | 13226.523 | 11112.051 |
| Japan | EG.1 | 1.057 | 1.040 | 1.074 | 1.000 | 11895.541 | 11387.662 |
| Japan | FL.4 | 1.056 | 1.042 | 1.071 | 1.000 | 10846.646 | 11131.328 |
| Japan | FU.1 | 1.054 | 1.029 | 1.079 | 1.000 | 16282.693 | 11357.217 |
| Japan | XBB.1.16 | 1.052 | 1.041 | 1.063 | 1.000 | 7094.965 | 9648.008 |
| Japan | EG.2 | 1.050 | 1.024 | 1.076 | 1.000 | 17794.055 | 12395.230 |
| Japan | CJ.1.3 | 1.041 | 1.013 | 1.071 | 1.000 | 19324.697 | 12612.005 |
| Japan | XBB.2.3 | 1.036 | 1.013 | 1.059 | 1.000 | 16691.648 | 12163.123 |
| Japan | XBB.1.16.1 | 1.035 | 1.022 | 1.049 | 1.000 | 10098.495 | 11788.071 |
| Japan | XBB.1.9.2 | 1.034 | 1.013 | 1.055 | 1.000 | 14843.659 | 11463.308 |
| Japan | FL.5 | 1.031 | 1.010 | 1.053 | 1.000 | 15482.051 | 11078.203 |
| Japan | XBB.1.9.1 | 1.015 | 1.002 | 1.028 | 1.000 | 8649.161 | 10967.809 |
| Japan | FL.2 | 1.004 | 0.986 | 1.023 | 1.001 | 14069.411 | 11517.641 |
| Japan | XBB.1.5.24 | 1.002 | 0.976 | 1.026 | 1.000 | 19750.344 | 12504.687 |
| Japan | GR.1 | 0.996 | 0.969 | 1.023 | 1.001 | 18199.213 | 11259.637 |
| Japan | FD.3 | 0.996 | 0.961 | 1.029 | 1.000 | 22137.245 | 12226.277 |
| Japan | EG.4 | 0.989 | 0.956 | 1.021 | 1.000 | 21166.280 | 12451.982 |
| Japan | FR.1 | 0.983 | 0.956 | 1.010 | 1.000 | 19909.113 | 12160.149 |
| Japan | CK.1.1 | 0.970 | 0.933 | 1.007 | 1.000 | 21290.210 | 10780.000 |
| Japan | XBB.1.5.1 | 0.959 | 0.926 | 0.991 | 1.000 | 20518.867 | 12622.277 |
| Japan | BN.1.2 | 0.955 | 0.927 | 0.983 | 1.000 | 19839.424 | 12244.441 |
| Japan | BF.7.15 | 0.951 | 0.931 | 0.970 | 1.000 | 15615.758 | 12124.496 |
| Japan | BQ.1.1.76 | 0.945 | 0.906 | 0.982 | 1.000 | 22460.318 | 12007.958 |
| Japan | BQ.1.1.45 | 0.928 | 0.884 | 0.970 | 1.001 | 24045.347 | 12339.500 |
| Japan | BF.7.4.1 | 0.927 | 0.896 | 0.957 | 1.001 | 21116.280 | 11736.014 |
| Japan | BN.1.3.2 | 0.921 | 0.880 | 0.960 | 1.000 | 23324.227 | 11999.601 |
| Japan | BF.11 | 0.917 | 0.876 | 0.955 | 1.001 | 24269.762 | 12790.234 |
| Japan | BQ.1.1 | 0.903 | 0.872 | 0.933 | 1.000 | 21117.241 | 11793.493 |
| Japan | BN.1.3 | 0.889 | 0.844 | 0.932 | 1.000 | 25142.002 | 11362.317 |
| Japan | XBB.1.5.23 | 0.889 | 0.842 | 0.933 | 1.000 | 23604.501 | 12450.849 |
| Singapore | EG.5.1 | 1.235 | 1.180 | 1.297 | 1.000 | 8132.463 | 9007.638 |
| Singapore | XBB.2.3.11 | 1.059 | 1.036 | 1.084 | 1.001 | 5005.086 | 8520.465 |
| Singapore | FU.1 | 1.058 | 1.039 | 1.077 | 1.000 | 3509.181 | 6040.722 |
| Singapore | XBB.1.16.2 | 1.048 | 1.027 | 1.070 | 1.000 | 4189.416 | 6994.983 |
| Singapore | EG.2 | 1.040 | 1.018 | 1.063 | 1.000 | 4365.164 | 6979.150 |
| Singapore | XBB.1.16.1 | 1.032 | 1.017 | 1.048 | 1.001 | 2775.862 | 5366.814 |
| Singapore | XBB.1.16 | 1.027 | 1.012 | 1.043 | 1.001 | 2861.123 | 5464.513 |
| Singapore | XBB.2.3.2 | 1.020 | 1.005 | 1.035 | 1.001 | 2804.818 | 5384.690 |
| Singapore | FL.4 | 1.013 | 0.995 | 1.031 | 1.000 | 3291.625 | 6193.186 |
| Singapore | XBB.1.16.7 | 0.995 | 0.970 | 1.021 | 1.000 | 5638.694 | 9200.828 |
| Singapore | XBB.2.3 | 0.993 | 0.967 | 1.018 | 1.000 | 5444.831 | 8987.494 |
| Singapore | XBB.1.9.2 | 0.985 | 0.964 | 1.005 | 1.000 | 4659.137 | 8341.666 |
| Singapore | XBB.1.9.1 | 0.983 | 0.963 | 1.003 | 1.000 | 4261.377 | 7804.220 |
| Singapore | XBB.2.3.6 | 0.983 | 0.965 | 1.000 | 1.001 | 3693.712 | 6641.019 |
| Singapore | XBB.1.22.2 | 0.981 | 0.955 | 1.007 | 1.000 | 5908.600 | 9171.395 |
| Singapore | FP.1 | 0.962 | 0.941 | 0.984 | 1.000 | 5181.022 | 8653.313 |
| South Korea | GJ.1 | 1.204 | 1.157 | 1.254 | 1.001 | 26821.178 | 11252.485 |
| South Korea | EG.5.1 | 1.187 | 1.165 | 1.210 | 1.000 | 22425.723 | 11541.056 |
| South Korea | FU.1 | 1.160 | 1.130 | 1.192 | 1.000 | 25344.324 | 10380.414 |
| South Korea | FL.15 | 1.142 | 1.111 | 1.175 | 1.001 | 27156.528 | 11454.385 |
| South Korea | FL.10.1 | 1.132 | 1.111 | 1.154 | 1.000 | 22807.406 | 11980.816 |
| South Korea | XBB.2.3.8 | 1.102 | 1.092 | 1.113 | 1.000 | 14007.153 | 12165.577 |
| South Korea | FL.14 | 1.099 | 1.069 | 1.131 | 1.000 | 26322.893 | 11281.279 |
| South Korea | EG.5.2 | 1.097 | 1.066 | 1.128 | 1.000 | 25550.185 | 10861.972 |
| South Korea | FK.1.1 | 1.095 | 1.074 | 1.118 | 1.000 | 21555.860 | 11470.359 |
| South Korea | EG.2 | 1.095 | 1.064 | 1.127 | 1.001 | 24973.929 | 10184.185 |
| South Korea | XBB.2.3.2 | 1.095 | 1.077 | 1.113 | 1.000 | 21059.013 | 11593.645 |
| South Korea | XBB.1.16.2 | 1.092 | 1.078 | 1.107 | 1.001 | 19951.152 | 11903.790 |
| South Korea | EG.1.4 | 1.090 | 1.073 | 1.107 | 1.000 | 20969.523 | 11093.139 |
| South Korea | FL.2 | 1.087 | 1.069 | 1.105 | 1.000 | 18725.661 | 11577.360 |
| South Korea | XBB.1.16 | 1.083 | 1.074 | 1.091 | 1.000 | 12323.214 | 12067.968 |
| South Korea | FE.1.1 | 1.082 | 1.057 | 1.108 | 1.000 | 23801.663 | 11309.631 |
| South Korea | FL.7 | 1.074 | 1.046 | 1.102 | 1.000 | 27402.526 | 11896.497 |
| South Korea | XBB.2.3.6 | 1.073 | 1.045 | 1.102 | 1.000 | 24792.121 | 11770.817 |
| South Korea | XBB.2.3 | 1.072 | 1.056 | 1.087 | 1.000 | 19115.849 | 12182.297 |
| South Korea | FL.1.2 | 1.068 | 1.043 | 1.094 | 1.000 | 24303.272 | 12505.709 |
| South Korea | XBB.1.16.1 | 1.065 | 1.051 | 1.079 | 1.000 | 19027.002 | 12664.928 |
| South Korea | XBB.1.16.10 | 1.063 | 1.046 | 1.080 | 1.001 | 20740.307 | 11781.284 |
| South Korea | EG.1 | 1.061 | 1.053 | 1.068 | 1.000 | 10849.565 | 12003.285 |
| South Korea | FL.2.1 | 1.054 | 1.032 | 1.077 | 1.000 | 24379.259 | 11720.984 |
| South Korea | FL.1.3 | 1.053 | 1.038 | 1.069 | 1.000 | 19844.171 | 12602.915 |
| South Korea | FL.4 | 1.052 | 1.041 | 1.063 | 1.000 | 15237.541 | 11733.625 |
| South Korea | XBB.1.9.2 | 1.050 | 1.041 | 1.060 | 1.000 | 13088.159 | 12901.111 |
| South Korea | XBB.1.9.1 | 1.049 | 1.042 | 1.057 | 1.000 | 11021.529 | 11977.921 |
| South Korea | EG.4 | 1.045 | 1.019 | 1.072 | 1.000 | 25359.738 | 11982.837 |
| South Korea | XBB.1.5.49 | 1.041 | 1.016 | 1.067 | 1.001 | 24713.560 | 11558.151 |
| South Korea | GR.1 | 1.041 | 1.024 | 1.058 | 1.000 | 22610.673 | 13021.821 |
| South Korea | FL.5 | 1.040 | 1.021 | 1.059 | 1.000 | 20444.998 | 12604.697 |
| South Korea | FL.17.1 | 1.017 | 0.990 | 1.045 | 1.000 | 24638.318 | 11459.372 |
| South Korea | XBB.1.5.23 | 1.009 | 0.988 | 1.030 | 1.000 | 21815.671 | 12427.124 |
| South Korea | FL.13 | 1.007 | 0.985 | 1.028 | 1.000 | 24414.314 | 11964.621 |
| South Korea | FP.4 | 1.004 | 0.979 | 1.028 | 1.001 | 22520.137 | 12601.265 |
| South Korea | XBB.1.5.24 | 0.999 | 0.979 | 1.020 | 1.000 | 23463.246 | 12181.593 |
| South Korea | XBB.1.5.64 | 0.999 | 0.976 | 1.022 | 1.000 | 22986.751 | 12435.895 |
| South Korea | XBB.1.5.17 | 0.977 | 0.959 | 0.996 | 1.000 | 23898.131 | 12323.604 |
| South Korea | BN.1.2.2 | 0.972 | 0.946 | 0.999 | 1.000 | 23930.357 | 11804.581 |
| South Korea | BN.1.2.3 | 0.941 | 0.916 | 0.966 | 1.001 | 26065.860 | 11405.366 |
| South Korea | BQ.1.24 | 0.936 | 0.915 | 0.956 | 1.000 | 25220.749 | 12631.096 |
| South Korea | CH.1.1 | 0.931 | 0.913 | 0.949 | 1.000 | 23012.674 | 12195.537 |
| South Korea | CJ.1.3 | 0.928 | 0.914 | 0.942 | 1.001 | 21110.705 | 12443.726 |
| South Korea | BN.1.3 | 0.920 | 0.904 | 0.936 | 1.000 | 21246.728 | 12592.619 |
| South Korea | BN.1.2 | 0.913 | 0.897 | 0.929 | 1.000 | 22503.566 | 12255.534 |
| South Korea | BN.1.3.5 | 0.908 | 0.873 | 0.941 | 1.000 | 28259.078 | 12208.469 |
| South Korea | BN.1.2.5 | 0.901 | 0.879 | 0.923 | 1.000 | 25348.568 | 12621.264 |
| South Korea | FR.2 | 0.875 | 0.844 | 0.905 | 1.001 | 25956.062 | 12361.490 |
| USA | XBB.1.16.6 | 1.201 | 1.175 | 1.228 | 1.000 | 16410.712 | 12536.031 |
| USA | EG.5.1 | 1.196 | 1.179 | 1.214 | 1.001 | 18918.558 | 12329.176 |
| USA | GJ.1 | 1.194 | 1.169 | 1.220 | 1.001 | 17013.484 | 11778.098 |
| USA | EG.6.1 | 1.169 | 1.136 | 1.203 | 1.000 | 16673.797 | 11531.235 |
| USA | FD.1.1 | 1.161 | 1.131 | 1.193 | 1.000 | 18345.408 | 11955.003 |
| USA | XBB.1.5.72 | 1.159 | 1.135 | 1.184 | 1.001 | 18211.762 | 11834.063 |
| USA | EG.5.2 | 1.129 | 1.110 | 1.149 | 1.000 | 18136.499 | 11953.533 |
| USA | FU.2 | 1.128 | 1.101 | 1.155 | 1.001 | 18605.981 | 11313.770 |
| USA | FY.2 | 1.121 | 1.093 | 1.151 | 1.000 | 19137.312 | 11230.735 |
| USA | FU.1 | 1.120 | 1.105 | 1.134 | 1.001 | 18484.296 | 11676.712 |
| USA | FE.1.1 | 1.117 | 1.091 | 1.144 | 1.000 | 17348.747 | 12275.666 |
| USA | XBB.2.3.2 | 1.113 | 1.100 | 1.127 | 1.000 | 18213.638 | 12405.565 |
| USA | XBB.2.3.3 | 1.109 | 1.089 | 1.130 | 1.001 | 17532.282 | 12311.823 |
| USA | XBB.1.16.2 | 1.108 | 1.095 | 1.122 | 1.000 | 18615.970 | 12800.675 |
| USA | XBB.1.16.1 | 1.100 | 1.092 | 1.108 | 1.000 | 16267.451 | 12673.873 |
| USA | EG.1 | 1.097 | 1.086 | 1.109 | 1.000 | 17459.058 | 11702.997 |
| USA | GR.1 | 1.097 | 1.071 | 1.123 | 1.000 | 19651.901 | 11892.997 |
| USA | XBB.2.3 | 1.096 | 1.085 | 1.107 | 1.000 | 18266.846 | 12320.105 |
| USA | XBB.1.22 | 1.093 | 1.072 | 1.115 | 1.000 | 17460.540 | 12538.347 |
| USA | XBB.1.5.77 | 1.092 | 1.075 | 1.108 | 1.000 | 16383.505 | 12288.488 |
| USA | EG.2 | 1.089 | 1.062 | 1.117 | 1.000 | 18241.409 | 12223.474 |
| USA | XBB.1.16 | 1.088 | 1.082 | 1.093 | 1.001 | 14473.584 | 12779.955 |
| USA | XBB.1.5.44 | 1.084 | 1.061 | 1.107 | 1.000 | 17642.435 | 11739.137 |
| USA | XBB.1.5.59 | 1.084 | 1.057 | 1.111 | 1.000 | 18699.869 | 12152.325 |
| USA | XBB.1.5.28 | 1.082 | 1.067 | 1.097 | 1.000 | 18618.327 | 12022.392 |
| USA | FL.4 | 1.081 | 1.070 | 1.092 | 1.001 | 17930.002 | 12133.524 |
| USA | FE.1.2 | 1.080 | 1.060 | 1.101 | 1.000 | 18770.429 | 11970.842 |
| USA | FL.3 | 1.080 | 1.056 | 1.104 | 1.000 | 20246.380 | 12678.376 |
| USA | XBB.1.5.68 | 1.077 | 1.062 | 1.092 | 1.000 | 19759.163 | 12580.766 |
| USA | XBB.1.5.37 | 1.075 | 1.058 | 1.093 | 1.000 | 18145.562 | 12607.805 |
| USA | EU.1.1 | 1.074 | 1.057 | 1.090 | 1.000 | 18444.647 | 12263.702 |
| USA | EG.1.4 | 1.072 | 1.048 | 1.097 | 1.000 | 19430.973 | 12725.575 |
| USA | XBB.1.5.10 | 1.065 | 1.054 | 1.076 | 1.000 | 16379.942 | 12538.586 |
| USA | XBB.1.5.39 | 1.063 | 1.036 | 1.091 | 1.001 | 18204.317 | 12804.718 |
| USA | FL.2 | 1.062 | 1.048 | 1.076 | 1.000 | 16842.660 | 12681.152 |
| USA | XBB.1.9.1 | 1.062 | 1.055 | 1.069 | 1.000 | 18235.379 | 13280.757 |
| USA | FL.13 | 1.057 | 1.032 | 1.082 | 1.000 | 18943.728 | 12995.630 |
| USA | XBB.1.5.86 | 1.055 | 1.029 | 1.080 | 1.000 | 18968.214 | 12150.805 |
| USA | XBB.1.5.73 | 1.054 | 1.034 | 1.073 | 1.000 | 18520.659 | 13017.283 |
| USA | FL.5 | 1.049 | 1.032 | 1.066 | 1.000 | 18018.316 | 12742.318 |
| USA | XBB.1.9.2 | 1.041 | 1.030 | 1.051 | 1.000 | 17988.209 | 12412.351 |
| USA | XBB.1.19.1 | 1.033 | 1.009 | 1.057 | 1.000 | 18484.396 | 12887.096 |
| USA | XBB.1.17.1 | 1.033 | 1.005 | 1.061 | 1.000 | 18716.936 | 12167.809 |
| USA | XBB.1.5.18 | 1.032 | 1.003 | 1.061 | 1.000 | 17260.252 | 12056.514 |
| USA | XBB.1.5.56 | 1.031 | 1.004 | 1.058 | 1.000 | 18807.117 | 12600.717 |
| USA | XBB.1.5.12 | 1.030 | 1.005 | 1.054 | 1.000 | 17995.093 | 12958.361 |
| USA | XBB.1.5.80 | 1.029 | 1.002 | 1.055 | 1.000 | 16127.215 | 13053.516 |
| USA | XBB.1.5.30 | 1.025 | 1.007 | 1.043 | 1.000 | 17869.061 | 11544.140 |
| USA | XBB.1.5.27 | 1.024 | 1.003 | 1.046 | 1.000 | 19764.876 | 12810.216 |
| USA | EG.4 | 1.024 | 1.005 | 1.044 | 1.000 | 17505.779 | 11955.009 |
| USA | XBB.1.5.48 | 1.024 | 1.007 | 1.041 | 1.000 | 17092.683 | 12631.377 |
| USA | XBB.1.5.50 | 1.024 | 0.997 | 1.050 | 1.000 | 19099.514 | 12640.215 |
| USA | FL.12 | 1.023 | 0.994 | 1.051 | 1.001 | 19924.466 | 12717.493 |
| USA | XBB.1.5.49 | 1.020 | 1.013 | 1.028 | 1.000 | 18260.512 | 12379.533 |
| USA | XBB.1.5.1 | 1.020 | 1.010 | 1.029 | 1.000 | 18389.446 | 12267.234 |
| USA | XBB.1.5.24 | 1.019 | 0.997 | 1.041 | 1.000 | 19023.282 | 13218.202 |
| USA | XBB.1.5.4 | 1.018 | 1.004 | 1.031 | 1.000 | 18461.555 | 12655.774 |
| USA | FL.10 | 1.018 | 0.992 | 1.043 | 1.000 | 18792.308 | 12915.634 |
| USA | XBB.1.5.3 | 1.012 | 0.986 | 1.038 | 1.000 | 19810.276 | 12893.577 |
| USA | XBB.1.5.35 | 1.012 | 0.998 | 1.026 | 1.000 | 18355.999 | 12100.719 |
| USA | XBB.1.5.17 | 1.011 | 0.999 | 1.023 | 1.000 | 17662.708 | 12242.314 |
| USA | XBB.1.5.61 | 1.008 | 0.980 | 1.035 | 1.001 | 19314.606 | 13238.328 |
| USA | XBB.1.5.16 | 1.006 | 0.987 | 1.025 | 1.000 | 19011.037 | 12860.989 |
| USA | XBB.1.5.20 | 1.005 | 0.988 | 1.022 | 1.000 | 19191.794 | 12571.352 |
| USA | XBB.1.5.31 | 1.004 | 0.983 | 1.025 | 1.000 | 19544.483 | 13065.938 |
| USA | XBB.1.5.14 | 1.004 | 0.975 | 1.031 | 1.000 | 19798.408 | 11947.783 |
| USA | XBB.1.5.15 | 1.002 | 0.991 | 1.013 | 1.000 | 18082.609 | 12591.057 |
| USA | FD.4 | 1.002 | 0.982 | 1.021 | 1.000 | 18758.790 | 12412.018 |
| USA | XBB.1.5.7 | 1.001 | 0.980 | 1.022 | 1.000 | 20528.201 | 12445.128 |
| USA | XBB.1.5.51 | 1.001 | 0.984 | 1.018 | 1.000 | 17757.650 | 14099.442 |
| USA | XBB.1.5.78 | 1.001 | 0.977 | 1.024 | 1.000 | 19564.041 | 13006.153 |
| USA | EK.2 | 1.000 | 0.969 | 1.030 | 1.000 | 19820.082 | 13051.270 |
| USA | CH.1.1 | 0.995 | 0.962 | 1.026 | 1.000 | 18244.659 | 13174.267 |
| USA | XBB.1.5.9 | 0.992 | 0.959 | 1.026 | 1.000 | 19099.569 | 12765.173 |
| USA | XBB.1.5.62 | 0.991 | 0.962 | 1.019 | 1.000 | 17983.481 | 12270.645 |
| USA | XBB.1.5.41 | 0.987 | 0.955 | 1.017 | 1.000 | 15573.811 | 12338.776 |
| USA | XBB.1.5.2 | 0.986 | 0.966 | 1.007 | 1.000 | 18766.951 | 11257.246 |
| USA | XBB.1.5.13 | 0.986 | 0.974 | 0.998 | 1.000 | 18391.317 | 12406.034 |
| USA | XBB.1.5.32 | 0.982 | 0.966 | 0.998 | 1.000 | 19360.960 | 13101.399 |
| USA | XBB.1.5.52 | 0.981 | 0.961 | 1.001 | 1.000 | 19273.741 | 13174.876 |
| USA | XBB.1.5.66 | 0.978 | 0.957 | 0.998 | 1.000 | 18534.671 | 13452.254 |
| USA | XBB.1.5.21 | 0.976 | 0.956 | 0.996 | 1.000 | 18110.950 | 12721.728 |
| USA | XBB.1.5.11 | 0.974 | 0.950 | 0.998 | 1.000 | 17570.825 | 13098.179 |
| USA | BQ.1.1 | 0.968 | 0.942 | 0.993 | 1.000 | 19096.719 | 12964.322 |
| USA | XBB.1.5.69 | 0.961 | 0.938 | 0.984 | 1.000 | 20636.413 | 12125.611 |
| USA | FD.2 | 0.957 | 0.943 | 0.970 | 1.000 | 20007.236 | 13440.453 |
| USA | XBB.1.5.33 | 0.947 | 0.926 | 0.967 | 1.000 | 19811.713 | 13486.054 |
| USA | XBB.1.5.19 | 0.947 | 0.916 | 0.976 | 1.000 | 17703.735 | 13347.167 |
| USA | XBB.1.5.5 | 0.936 | 0.899 | 0.970 | 1.000 | 17211.557 | 13037.697 |

The Re value of XBB.1.5 is set at 1.

**Table S3.**
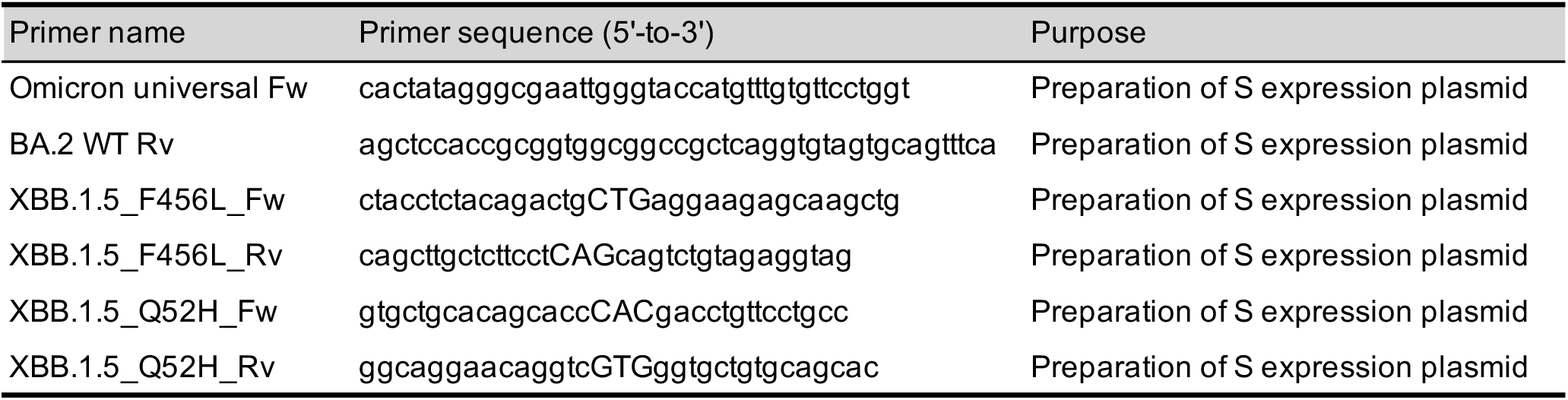
Primers used in this study.

**Figure S1.**
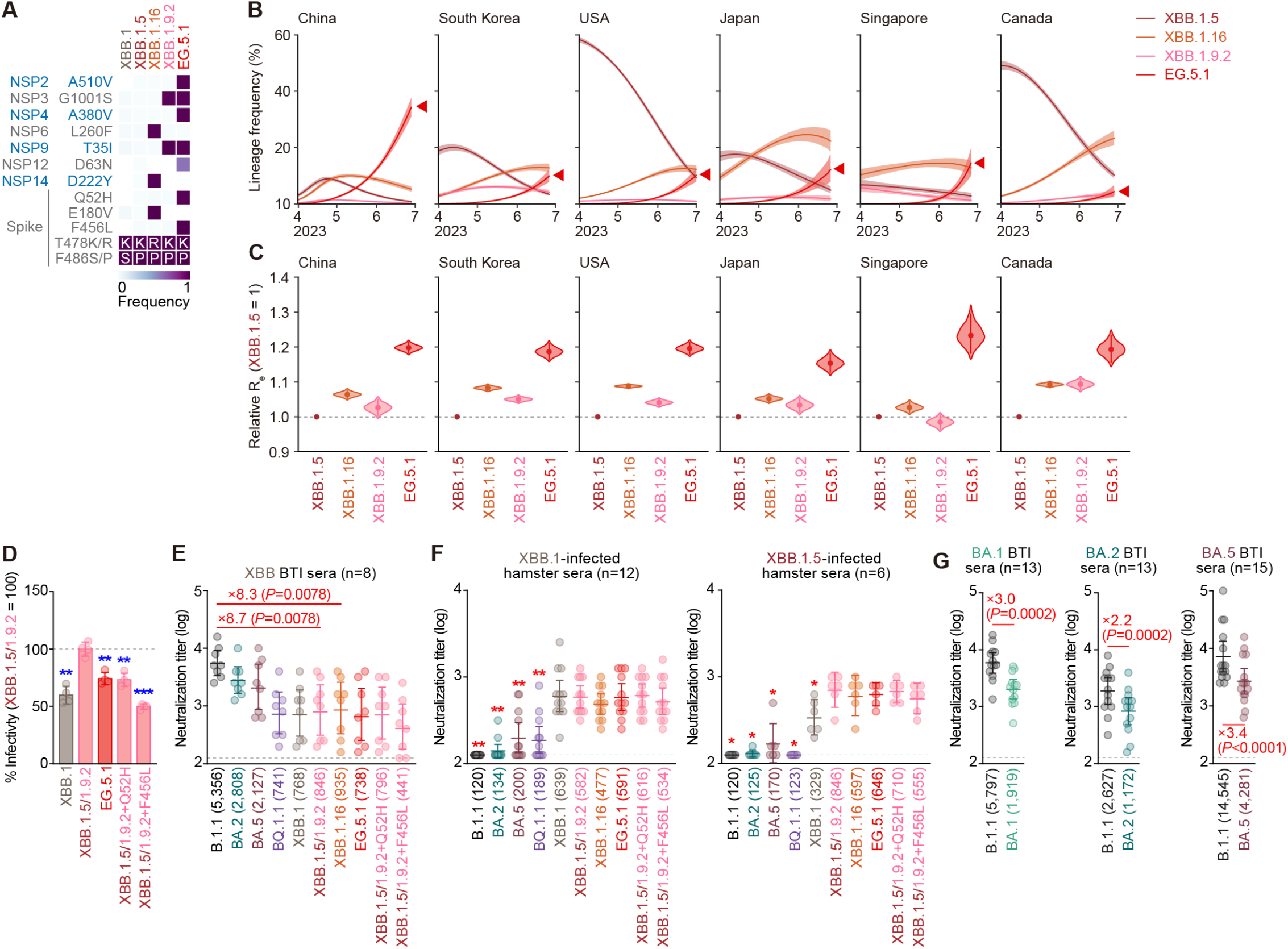
Virological features of EG.5.1 and XBB BTI. (**A**) Frequency of mutations of interest in the representative XBB sublineages. Only mutations with a frequency >0.5 in at least one but not all the representative sublineages are shown. Note that the S proteins of XBB.1.5 and XBB.1.9.2 are identical. (**B**) Estimated epidemic dynamics of the representative XBB sublineages in countries where >50 sequences of EG.5.1, XBB.1.5, XBB.1.9.2, and XBB.1.16 were detected from April 1, 2023 to July 13, 2023. Countries are ordered according to the number of detected sequences of EG.5.1. Line, posterior mean; ribbon, 95% Bayesian confidence interval. The dynamics for EG.5.1 is highlighted by a red arrowhead. (**C**) Estimated relative Re of the representative XBB sublineages in the six countries. The relative Re of XBB.1.5 is set to 1 (horizontal dashed line). Violin, posterior distribution; dot, posterior mean; line, 95% Bayesian confidence interval. (**D**) Lentivirus-based pseudovirus assay. HOS-ACE2-TMPRSS2 cells were infected with pseudoviruses bearing each S protein. The amount of input virus was normalized to the amount of HIV-1 p24 capsid protein. The percentage infectivity of XBB.1.5/1.9.2, XBB.1.5/1.9.2+Q52H, XBB.1.5/1.9.2+F456L, and EG.5.1 compared to that of XBB.1.5/1.9.2 are shown. The horizontal dash line indicates the mean value of the percentage infectivity of the XBB.1.5/1.9.2. Assays were performed in quadruplicate. The presented data are expressed as the average ± SD. Each dot indicates the result of an individual replicate. (**E-G**) Neutralization assay. Assays were performed with pseudoviruses harboring the S proteins of B.1.1, BA.1, BA.2, BA.5, BQ.1.1, XBB.1, XBB.1.5/1.9.2, XBB.1.16, EG.5.1, XBB.1.5/1.9.2+Q52H, and XBB.1.5/1.9.2+F456L. The following sera were used: convalescent sera from fully vaccinated individuals who had been infected with XBB.1.5 (one 3-dose vaccinated. 1 donor in total), XBB.1.9 (one 3-dose vaccinated donor, one 4-dose vaccinated donor and one 5-dose vaccinated donor. 3 donors in total), and XBB.1.16 (one 2-dose vaccinated donor, two 3-dose vaccinated donors, and one 4-dose vaccinated donor. 4 donors in total) (**E**); sera from hamster infected with XBB.1 (left) or XBB.1.5 (right) (**F**); and convalescent sera from fully vaccinated individuals who had been infected with BA.1 (thirteen 2-dose vaccinated, 13 donors in total) (left)^1^, BA.2 (nine 2-dose vaccinated and four 3-dose vaccinated donors. 13 donors in total) (middle)^19^, and BA.5 (one 2-dose vaccinated, thirteen 3-dose vaccinated donors, and one 4-dose vaccinated. 15 donors in total) (right)^19^ (**G**). Each dot indicates the result of an individual replicate. Assays for each serum sample were performed in triplicate to determine the 50% neutralization titer (NT50). Each dot represents one NT50 value, and the geometric mean and 95% confidence interval are shown. The number in parenthesis indicates the mean of NT50 values. The horizontal dash line indicates the detection limit (120-fold). In **D**, statistically significant differences (**, *P* < 0.001, ***, *P* < 0.0001) versus XBB.1.5/1.9.2 were determined by two-sided Student’s *t* tests. Blue asterisks indicate decreased percentage of infectivity. In **E and G**, statistically significant differences versus B.1.1 were determined by two-sided Wilcoxon signed-rank tests. The fold change between B.1.1 and the variant indicated is shown in red. Background information on the convalescent donors is summarized in **Table S1**. In **F**, statistically significant differences (*, *P* < 0.01, **, *P* < 0.001) between B.1.1 and XBB.1 (left) or XBB.1.5/1.9.2 (right) were determined by two-sided Wilcoxon signed-rank tests and indicated with asterisks. Red asterisks indicate decreased NT50s.

## Consortia

### The Genotype to Phenotype Japan (G2P-Japan) Consortium

#### The Institute of Medical Science, The University of Tokyo, Japan

Naoko Misawa, Ziyi Guo, Alfredo A. Hinay Jr., Arnon Plianchaisuk, Jarel Elgin M. Tolentino, Luo Chen, Shigeru Fujita, Lin Pan, Gustav Joas, Olivia Putri, Yoonjin Kim, Mai Suganami, Mika Chiba, Ryo Yoshimura, Kyoko Yasuda, Keiko Iida, Naomi Ohsumi, Adam P. Strange, Shiho Tanaka

#### Hokkaido University, Japan

Takasuke Fukuhara, Tomokazu Tamura, Rigel Suzuki, Saori Suzuki, Hayato Ito, Keita Matsuno, Hirofumi Sawa, Naganori Nao, Shinya Tanaka, Masumi Tsuda, Lei Wang, Yoshikata Oda, Zannatul Ferdous, Kenji Shishido

#### Tokai University, Japan

So Nakagawa

#### Kyoto University, Japan

Kotaro Shirakawa, Akifumi Takaori-Kondo, Kayoko Nagata, Ryosuke Nomura, Yoshihito Horisawa, Yusuke Tashiro, Yugo Kawai, Kazuo Takayama, Rina Hashimoto, Sayaka Deguchi, Yukio Watanabe, Ayaka Sakamoto, Naoko Yasuhara, Takao Hashiguchi, Tateki Suzuki, Kanako Kimura, Jiei Sasaki, Yukari Nakajima, Hisano Yajima

#### Hiroshima University, Japan

Takashi Irie, Ryoko Kawabata

#### Kyushu University, Japan

Kaori Tabata

#### Kumamoto University, Japan

Terumasa Ikeda, Hesham Nasser, Ryo Shimizu, MST Monira Begum, Michael Jonathan, Yuka Mugita, Otowa Takahashi, Kimiko Ichihara, Takamasa Ueno, Chihiro Motozono, Mako Toyoda

#### University of Miyazaki, Japan

Akatsuki Saito, Maya Shofa, Yuki Shibatani, Tomoko Nishiuchi

##### Acknowledgments

We would like to thank all members of The Genotype to Phenotype Japan (G2P-Japan) Consortium. We thank Dr. Kenzo Tokunaga (National Institute of Infectious Diseases, Japan) for sharing materials. We gratefully acknowledge the numerous laboratories worldwide that have provided sequence data and metadata to GISAID. A full list of originating and submitting laboratories for the sequences used in our analysis can be found at https://www.gisaid.org using the EPI-SET-ID: EPI_SET_230725pv.

